# A cellular assay to determine the fusion capacity of MFN2 variants linked to Charcot-Marie-Tooth type 2A

**DOI:** 10.1101/2024.03.11.584414

**Authors:** Chloe Barsa, Julian Perrin, Claudine David, Arnaud Mourier, Manuel Rojo

## Abstract

Charcot-Marie-Tooth Disease (CMT) is an inherited peripheral neuropathy with two main forms: demyelinating CMT1 and axonal CMT2. The most frequent subtype of CMT2 (CMT2A) is linked to mutations of *MFN2*, encoding a membrane-anchored GTP-binding protein essential for mitochondrial outer membrane fusion. The use of Next-Generation Sequencing for genetic analysis has led to the identification of increasing numbers of *MFN2* variants, but a majority of them remain variants of unknown significance, depriving patients of a clear diagnosis. In this work, we establish a cellular assay allowing to assess the impact of *MFN2* variants linked to CMT2A on the fusion capacity of MFN2. The analysis of 12 *MFN2* variants revealed that five abolish fusion, one induces an important reduction and six retain a fusion capacity similar to that of wild-type MFN2. Their analysis with computational variant effect predictors demonstrated a remarkable correlation of our results with predictions based on protein sequence analysis. This work develops novel tools to determine the functional impact of known and novel *MFN2* variants and identifies computational tools allowing to predict their possible consequences and pathogenicity.

## Introduction

Charcot–Marie–Tooth disease (CMT) is a hereditary sensory and motor neuropathy which belongs to a heterogeneous group of inherited peripheral neuropathies that includes peripheral sensitive and motor neuropathies. With autosomal dominant, autosomal recessive, and X linked inheritance, and an estimated prevalence of 1:2500 to 1:10000, CMTs are among the most frequently diagnosed hereditary neuropathies (Bacquet et al., 2018) (Pipis et al., 2019). Autosomal dominant CMTs exist in two main forms, demyelinating CMT1 and axonal CMT2 and the most prevalent symptoms (distal motor and sensory weakness) start to manifest in childhood or adolescence, but can also appear during adulthood (Choi et al., 2015; Pipis et al., 2020). Progress in molecular diagnosis of CMTs and related neuropathies revealed that some of them were linked to mutations in genes governing mitochondrial bioenergetics or dynamics (Bertholet et al., 2016; Bacquet et al., 2018; Pipis et al., 2019). The CMT of type 2A, the most frequent subtype of CMT2, is caused by mutations of *MFN2* (Züchner et al., 2004; Stuppia et al., 2015); it is mainly associated with autosomal dominant inheritance (Stuppia et al., 2015) (Pipis et al., 2020), but recessive and semi-dominant forms have been also described (Nicholson et al., 2008; Calvo et al., 2009; Tomaselli et al., 2016)

*MFN2* is a nuclear gene encoding a multifunctional protein (Filadi et al., 2018) that is essential for the fusion of the outer mitochondrial membrane (OMM) (Santel and Fuller, 2001; Rojo et al., 2002; Chen et al., 2003). MFN2 and its homologous protein MFN1 are ubiquitously expressed dynamin related proteins that coordinately regulate mitochondrial fusion (Chen et al., 2003). They are anchored to the cytosolic face of the OMM and contain a GTPase domain and two coiled-coil (or heptad repeat) domains separated by transmembrane domains (Figure 1A). MFN1 and MFN2 can mediate fusion separately, but previous works have demonstrated that MFN1 or MFN2 can also interact with each other to catalyze fusion in a cooperative, GTPase-dependent manner (Detmer and Chan, 2007). Despite significant progress in the biochemical and structural characterization of MFN1 and MFN2 (Daste et al., 2018; Yan et al., 2018; Li et al., 2019) and beyond the consensus that MFN1 and MFN2 can physically interact (Detmer and Chan, 2007), the precise molecular mechanisms and conformational changes involved in MFN mediated membrane fusion are still debated (Daumke and Roux, 2017; Cohen and Tareste, 2018). Intriguingly, despite their strong sequence homology and redundant activity, no disease or neuropathy has been linked to *MFN1* mutations.

**Figure 1.**
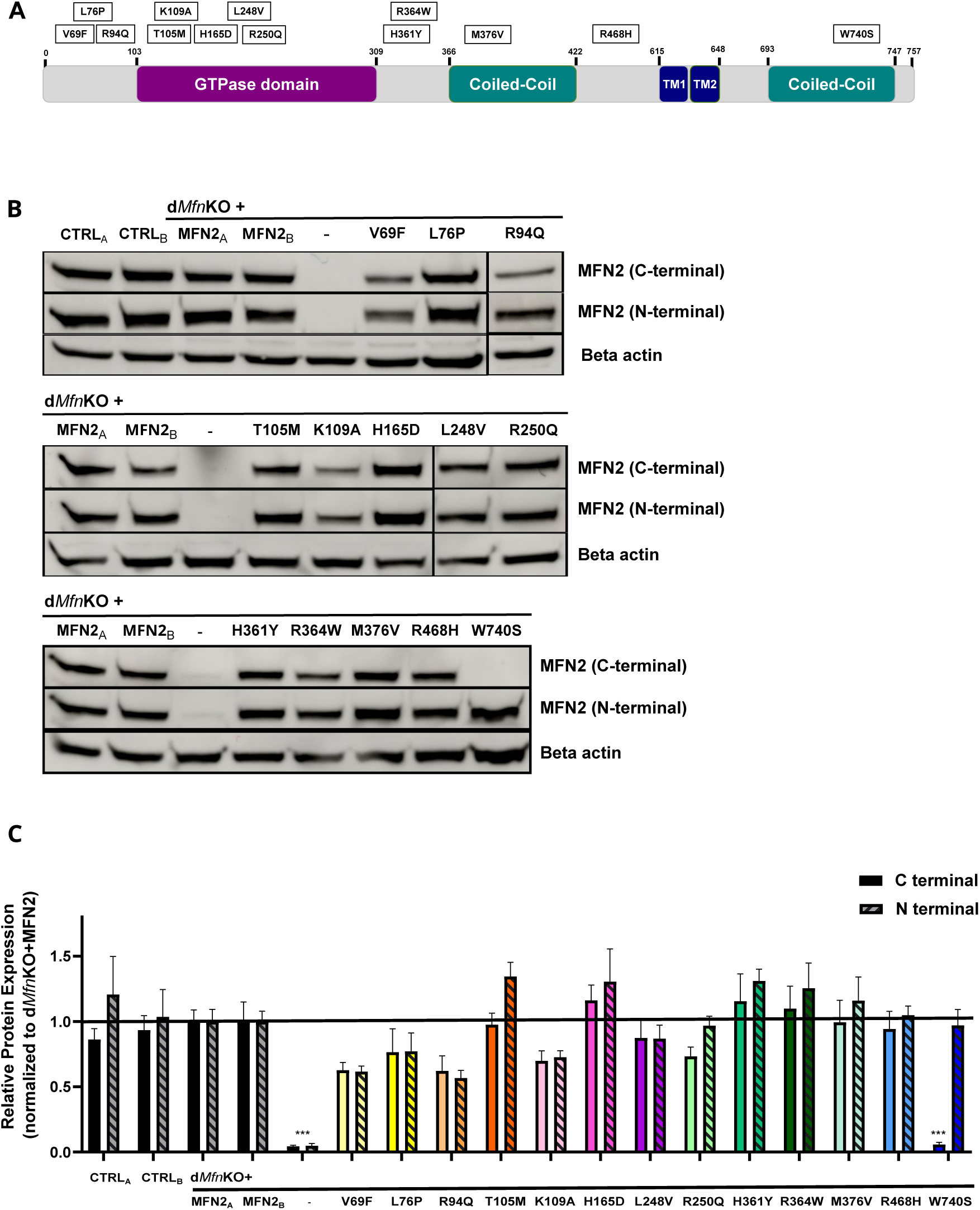
Expression of *MFN2* variants in double *Mfn* KO mouse embryo fibroblasts (d*Mfn*KO MEFs). **A**. Schematic illustration of the domain organization of MFN2 indicating the position of the SNVs characterized in this study, the position of the first and last amino acid of the MFN2 molecule or domain is indicated. TM: transmembrane domain. Rectangles indicate the amino acid change induced by SNVs and their relative position. **B**. Representative Western blots of MFN2 (mouse and rabbit antibodies targeting the C-terminal and N-terminal domain, respectively) and of a loading control (Beta actin) in different MEF lines. CTRL_A_, CTRL_B_: two different control MEF lines. MFN2_A_, MFN2_B_: two independently transduced d*Mfn*KO MEF lines expressing wild-type MFN2. **C**. Bar-graphs of the quantitative analysis of MFN2 relative protein levels. Blots of four independent experiments were analyzed and normalized to beta actin and to dMfnKO+MFN2 lines. Means ± SEM are plotted and a one-way ANOVA with Kruskal-Wallis Multiple Comparison post hoc test was run. *P < 0.05, **P < 0.01, ***P < 0.01 compared with the d*Mfn*KO+MFN2 MEF lines.

Since the *princeps* publication identifying *MFN2* as the main genetic factor responsible for CMT2A (Züchner et al., 2004), more than 100 *MFN2* variants have been detected in CMT2A patients (Stuppia et al., 2015). Mutations affect mainly the coding region and are most commonly missense; they concentrate in the highly conserved GTPase and coiled-coil domains, but can be found all across the MFN2 protein sequence (Stuppia et al., 2015; Pipis et al., 2020).

The widespread adoption of Next-Generation Sequencing (NGS) as a standard for genetic diagnosis has tremendously increased the number of known *MFN2* single nucleotide variants (SNVs). The number of *MFN2* variants of unknown significance (VUS) is constantly increasing and a large number of patients carrying *MFN2* variants cannot be provided with a clear diagnosis. The interpretation and classification of SNVs pathogenicity with established guidelines (Richards et al., 2015) most commonly relies on a balanced and critical analysis of several factors, including familial segregation, functional characterization of the proteins encoded by SNVs and analysis with a variety of bioinformatics tools for Variant Effect Prediction (VEP). However, the correct interpretation of functional studies and of computational predictions is a complex endeavor that requires data curation, assay interpretation, and software calibration (Richards et al., 2015; Brnich et al., 2019) (Pejaver et al., 2022).

The gap of knowledge in our understanding of pathogenic mechanisms underpinning CMT2A disorder has prompted scientists to study the impact of *MFN2* variants with numerous approaches (Zaman and Shutt, 2022), but the use of different biological materials and experimental systems – ranging from muscle or sural nerve biopsies to cultured skin fibroblasts – prevents a direct and faithful comparison between the different *MFN2* variants. Analyses performed on patients’ fibroblasts revealed that, with a notable exception (Rouzier et al., 2012), mitochondrial fusion and morphology appear unaffected (Loiseau et al., 2007) (Amiott et al., 2008) (Larrea et al., 2019), precluding the identification of phenotypic hallmarks of *MFN2* variants in this system.

So far, only a small subset of the known *MFN2* mutations have been characterized by expression in biological models with isogenic backgrounds. *MFN2* variants expressed in rodents (De Gioia et al., 2020; Strickland et al., 2014; Sato-Yamada et al., 2022) or in mouse embryonic fibroblasts (MEFs) have demonstrated the pathogenic nature of some *MFN2* SNVs, but also reported that numerous *MFN2* variants do not affect the fusion activity of MFN2 (Detmer and Chan, 2007). The limited number of MFN2 variants with experimentally determined functional impact or pathogenicity, alongside the increasing pressure from diagnostic centers to characterize *MFN2* VUS, prompted us to develop a strategy allowing a direct, straightforward, and unbiased functional characterization of human mutated MFN2.

In this study, we developed a cell-based assay allowing a direct functional assessment of the impact of *MFN2* variants on MFN2 driven mitochondrial fusion. This functional test expresses *MFN2* SNVs with high fidelity and controlled conditions in well-established and characterized double *Mfn1*/*Mfn2* knock-out mouse embryonic fibroblasts (d*Mfn*KO MEFs) (Koshiba et al., 2004; Chen et al., 2005; 2010; Fissi et al., 2018; Silva Ramos et al., 2019). This new functional test allowed the characterization of 12 SNVs of *MFN2* identified in CMT2A patients. Expression of SNVs with comparable levels in an isogenic genetic background allowed the identification of SNVs altering the fusion capacity of MFN2. Beyond providing a new classification of *MFN2* SNVs based on their functional consequences, the obtained results allowed a critical appraisal of *in silico* tools for ‘variant effect prediction (VEP)’ and the validation of a subset of tools predicting alterations of the fusion capacity with remarkable accuracy.

## Results

### Generation of isogenic cell disease models expressing human *MFN2* variants linked to CMT2A

To characterize the OMM-fusion capacity of *MFN2* SNVs, we generated stably transduced double *Mfn1*/*Mfn2* knock-out MEFs (d*Mfn*KO MEFs) unable to fuse the OMM and, consequently, mitochondria. We transduced human wild-type *MFN2*, a mutant (p.K109A) encoding fusion-incompetent MFN2 (Chen et al., 2003) as well as *MFN2* SNVs identified in CMT2A patients. In this study, we selected a series of 12 *MFN2* SNVs that have been detected in several families and/or patients (Table 1) and distribute to different regions and functional domains of MFN2 (Figure 1A). Ten of them are classified as pathogenic or likely pathogenic (Table 1) and three of them (p.R94Q, p.T105M, and p.H361Y) had their pathogenicity confirmed in transgenic rodent models recapitulating CMT-related symptoms (De Gioia et al., 2020; Sato-Yamada et al., 2022). We also selected two variants that present a high allele frequency in the gnomAD database (Table 1) and are classified as variants of unknown significance (p.R250Q and p.R468H) and/or likely benign (p.R468H). The selected *MFN2* SNVs containing ORF were generated by established mutagenesis procedures and transduced into d*Mfn*KO MEFs by retroviral transduction. The successful and faithful transduction of d*Mfn*KO MEFs was verified by amplification and sequencing of the genome-integrated *MFN2* SNVs cDNAs of the different stable MEFs lines generated.

**Table 1:** Identity, position, classification and frequency of *MFN2* variants characterized in this study. P: pathogenic, LP: likely pathogenic, CI: conflicting interpretations, VUS: variant of unknown significance. LB: likely benign. B: benign. (#): number of patients/families/entries. **cDNA**: the variants are described using the NM_014874.3 transcript reference sequence. **gnomAD**: The Genome Aggregation Database. **HGMD**: The Human Gene Mutation Database. **LOVD**: Leiden Open Variation Database. **INVB**: Inherited Neuropathy Variant Browser. Pipis et al. 2020. AD/AR/SD: autosomal dominant / autosomal recessive / semi-dominant inheritance.

|  |  | gnomAD v4.0.0 | HGMD | Clinvar | LOVD | INVB | Pipis et al. 2020 |  |
| --- | --- | --- | --- | --- | --- | --- | --- | --- |
| protein | cDNA | Frequency (Count) | Phenotype | Significance (#) | Classification (#) | (#) | Classification (Inheritance) | reference |
| p.V69F | c.205G>T | – | CMT2A | P (1) | P (2) | (1) | – | Züchner et al., 2004 |
| p.L76P | c.227T>C | 1.24e-6 (2) | CMT2A | P (6) | P (2) | (3) | LP (AD) | Züchner et al., 2004 |
| p.R94Q | c.281G>A | – | CMT2A | P (13) | P/VUS (5) | (8) | P (AD) | Züchner et al., 2004 |
| p.T105M | c.314C>T | – | CMT2A | P/LP (8) | P (3) | (7) | P (AD) | Kijima et al., 2005 |
| p.H165D | c.493C>G | – | CMT2A | P (3) | P (2) | (1) | – | Zhu et al., 2005 |
| p.L248V | c.742C>G | – | CMT2A | LP (2) | P (1) | (1) | LP (AD) | Feely et al., 2011 |
| p.R250Q | c.749G>A | 3.54e-4 (571) | CMT2A | CI (9) | P/VUS (4) | (2) | (AR / SD) | Verhoeven et al., 2006 |
| p.H361Y | c.1081C>T | – | CMT2A | P (2) | P (2) | (3) | P (AD) | Züchner et al., 2006 |
| p.R364W | c.1090C>T | – | CMT2A | P (16) | P (6) | (8) | P (AD) | Züchner et al., 2006 |
| p.M376V | c.1126A>G | – | CMT2A | P/LP (6) | P (4) | (2) | P (AD) | Casasnovas et al., 2010 |
| p.R468H | c.1403G>A | 2.76e-3 (4448) | CMT2A ? | CI (12) | P/LP/VUS/LB/B (18) | (3) | LB (AD) | Engelfried et al., 2006 |
| p.W740S | Nc.2219G>C | – | CMT2A | P/LP (13) | P (2) | (5) | P (AD) | Züchner et al., 2004 |

The genetic validation of the fifteen d*Mfn*KO MEFs lines, two expressing wild-type MFN2, one expressing fusion-incompetent p.K109A and 12 expressing *MFN2* variants linked to CMT2A, prompted us to validate by Western blot analysis that the MFN2 transgene is evenly expressed across these lines. To this end, total cell protein extracts were subjected to classical denaturing PAGE followed by Western blot analysis with two independent MFN2 antibodies targeting C-terminal or N-terminal epitopes and recognizing mouse and human MFN2 (Figure 1B and C). First, the almost identical levels of human MFN2 protein expressed in two independently generated stable d*Mfn*KO MEFs (MFN2_A_ and MFN2_B_), validated the high reproducibility and fidelity of our expression strategy (Figure 1B and C). Interestingly, the levels of human MFN2 proteins expressed in MFN2_A_ and MFN2_B_ was found to be almost identical to the levels of the endogenous MFN2 expressed in two control MEF lines (CTRL_A_ and CTRL_B_). The Western blot analyses performed with d*Mfn*KO MEFs expressing *MFN2* SNVs demonstrated that the levels of human MFN2 protein variants did not differ significantly from the human or murine MFN2 levels expressed in control MEFs (Figure 1B and C). Of note, the p.W740S variant was only detected with the antibody targeting the N-terminal domain (Figure 1B and C). The absence of signal with the C-terminal antibody is most probably explained by the fact that the amino acid change induced by p.W740S localizes to the epitope recognized by this monoclonal antibody. These results demonstrated (i) that our new approach can efficiently generate stable CMT2A disease model cells, evenly expressing MFN2 variants in an isogenic background, and (ii) that none of the analyzed mutations impact MFN2 expression or stability.

### Functional characterization of *MFN2* SNVs by quantitative analysis of MFN2-mediated mitochondrial fusion

The isogenic disease model selected to investigate the functional impact of CMT2A variants on MFN2 fusion activity is the well-established d*Mfn*KO MEF-line (Koshiba et al., 2004) (Chen et al., 2005) (Fissi et al., 2018) (Silva Ramos et al., 2019). The d*Mfn*KO MEF-model was chosen since, OMM-fusion being abolished, these cells present a completely fragmented mitochondrial network that allows to accurately follow the fusogenic activity of transgenically expressed MFN2 by following the restauration of a filamentous mitochondrial network (Detmer and Chan, 2007). The mitochondrial network morphology was visualized and quantified in paraformaldehyde fixed MEFs using state of the art immunostaining protocols with a well-established anti-VDAC antibody. The mitochondrial morphology was quantified by classification of ‘overall mitochondrial morphology’ in different categories (filamentous, intermediate or fragmented) in three independent experiments on at least 500 cells (Figure 2B, supplementary Figure 1). The analysis of epifluorescence images confirmed that, in stark contrast with the filamentous mitochondria observed in control MEFs (CTRL_A_ and CTRL_B_), d*Mfn*KO MEFs devoid of OMM-fusion display a completely fragmentated mitochondrial network (Figure 2, supplementary Figure 1) (Koshiba et al., 2004). Furthermore, quantitative analyses demonstrated that, in line with previous work (Fissi et al., 2018), the filamentous mitochondrial network morphology is almost completely restored upon expression of wild-type human MFN2 (MFN_A_ and MFN_B_). The almost identical level of mitochondrial network restoration observed between the independent MFN2_A_ and MFN2_B_ cell models evenly expressing MFN2 (Figure 1 A and B) nicely confirmed the reproducibility and robustness of our quantitative approach (Figure 2 A and B). We also generated d*Mfn*KO MEFs expressing a negative control: as expected, expression of the *MFN2*-K109A mutant known to abolish the GTPase and the fusogenic activity of MFN2 (Chen et al., 2003) did not rescue the fragmented mitochondrial morphology of OMM-fusion deficient d*Mfn*KO MEFs (Figure 2, supplementary Figure 1). This quantitative analysis demonstrated that our cell-based functional test could accurately discriminate between active and inactive MFN2.

**Figure 2.**
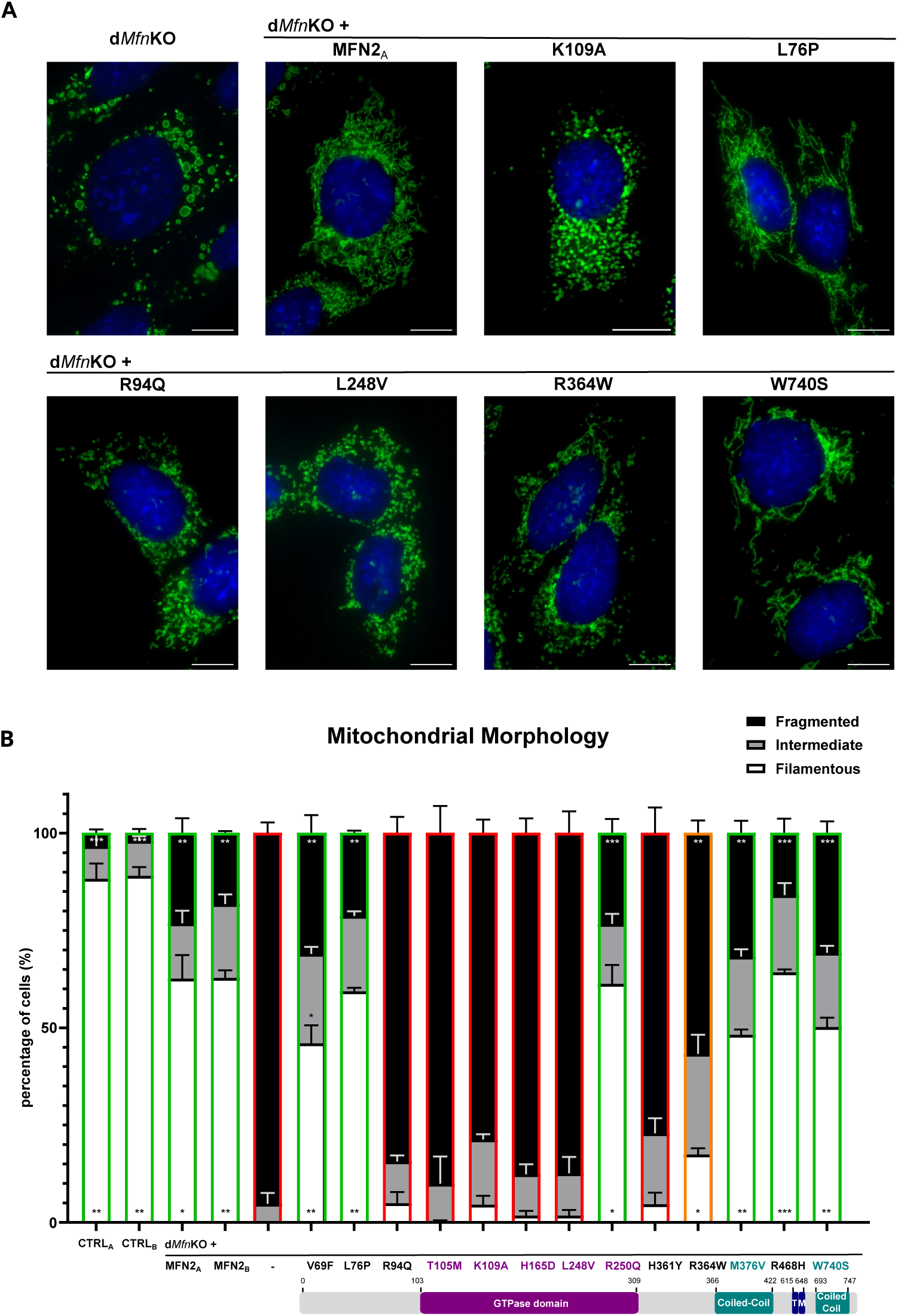
Visualization and quantitative analysis of mitochondrial morphology. **A.** Representative immunofluorescence images of untransduced d*Mfn*KO MEFs and of d*Mfn*KO MEFs expressing wild-type MFN2 (MFN2), a fusion incompetent mutant (K109A) or the indicated SNVs. MEFs were stained with the mitochondrial marker VDAC (green) and the nuclear stain DAPI (blue). Bar: 10 µm. **B.** Bar-graph of the quantitative analysis of mitochondrial morphology in ≥500 cells (3 independent experiments, 150-200 cells/experiment). The proportion of cells with filamentous, fragmented or intermediate mitochondrial morphology is expressed as % of cells. SNVs differ in their capacity to induce filament formation by fusion. The percentage of the different morphologies was compared to those of untransduced d*Mfn*KO cells using two-way ANOVA with Dunnett’s Multiple Comparison post hoc test. *P < 0.05, **P < 0.01, ***P < 0.01. SNVs differ in their capacity to restore fusion.

Next, we decided to challenge the cell-based functional assay by characterizing the 12 *MFN2* SNVs transduced into d*Mfn*KO. Our functional analyses revealed two main categories of MFN2 molecules (Figure 2, supplementary Figure 1): the group of SNVs that, like fusion-incompetent K109A, did not significantly rescue the aberrant morphology of d*Mfn*KO MEFs (p.R94Q, p.T105M, p.H165D, p.L248V and p.H361Y) and the group of SNVs that, alike wild-type MFN2, could efficiently restore the mitochondrial network (p.V69F, p.L76P, p.R250Q, p.M376V, p.R468H and p.W740S). A single variant, p.R364W, displayed an intermediate fusion phenotype with a capacity to restore OMM-fusion and a filamentous morphology that was detectable, but significantly affected in regard to the wild-type MFN2 (Figure 2). These results demonstrated that our cell-based OMM-fusion assay allows a robust quantification of mitochondrial morphology and the classification of human MFN2 SNVs according to their OMM-fusion capacity. Remarkably, our results reveal that roughly half of the SNVs identified in CMT2A patients were severely impairing MFN2’s fusion capacity.

### Subcellular localization of MFN2 variants reveals that SNVs do not affect mitochondrial targeting

To further characterize the impact of SNVs on MFN2 properties, we investigated whether SNVs provoke MFN2 mistargeting, a potential pathogenic mechanism accounting for MFN2 dysfunction. To this end, mitochondria were visualized with antibodies targeting TIM23, a subunit of the protein translocase located in the inner membrane, and the localization of MFN2 variants was determined by co-immunostaining with MFN2-specific antibodies. The specificity of the anti-MFN2 antibody was validated by the complete absence of signal in d*Mfn*KO MEFs (Figure 3, supplementary Figure 2), further supporting the fact that MFN2 detection was conditioned to transgenic expression of MFN2. Interestingly, these analyses unambiguously demonstrated that fusion competent as well as fusion-incompetent MFN2 mutants were properly targeted to mitochondria (Figure 3, supplementary Figure 2). These results demonstrated that none of the 12 SNVs characterized in this article alters mitochondrial targeting of MFN2 and that the impaired fusion capacity of some variants does not result from altered MFN2 localization.

**Figure 3.**
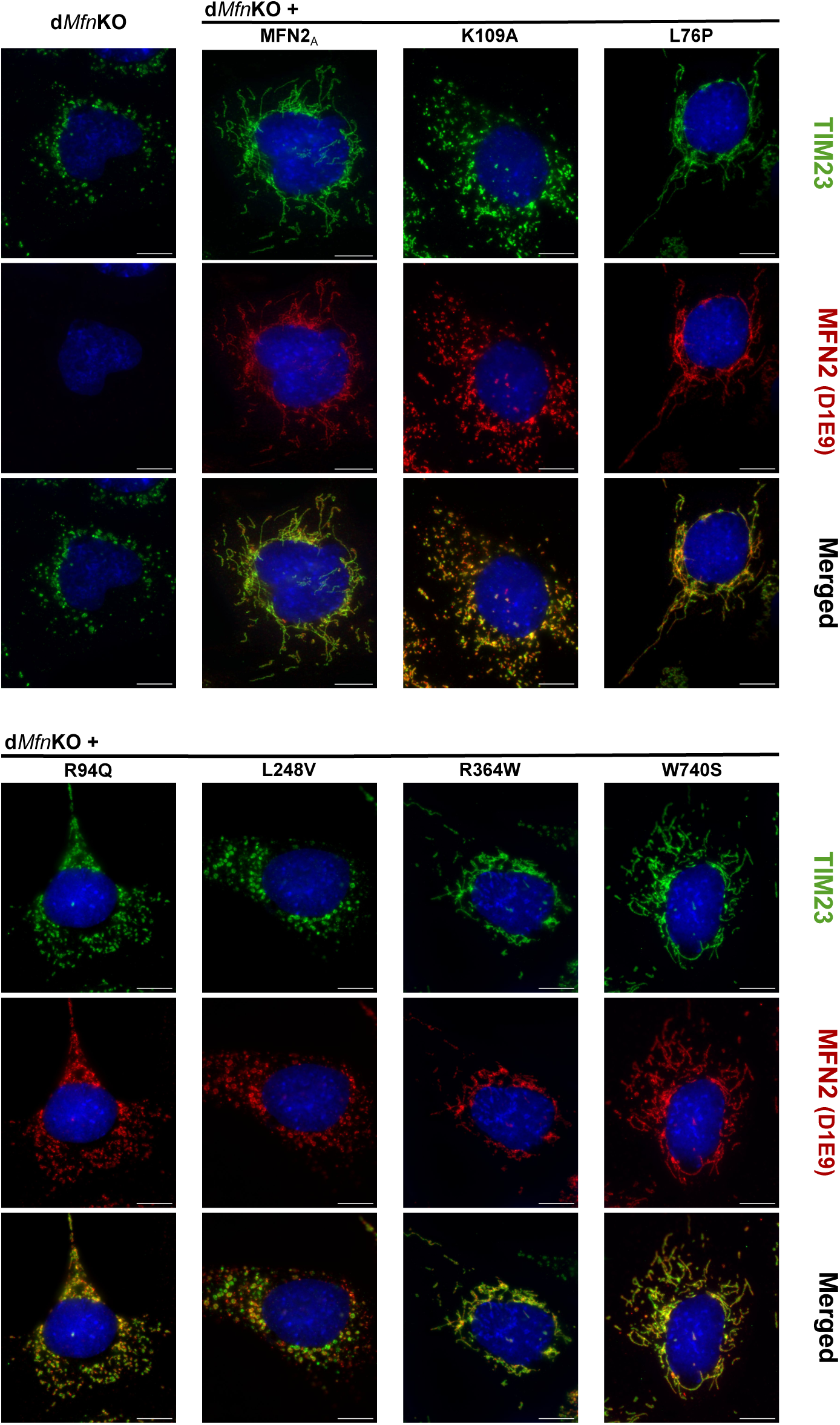
Mitochondrial localization of human MFN2 variants expressed in d*Mfn*KO MEFs. Representative immunofluorescence images of cells stained with antibodies against the mitochondrial marker TIM23 (green) and MFN2 (D1E9, red) and with the nuclear stain DAPI (blue). MFN2, undetectable in untransduced d*Mfn*KO MEFs, localizes to TIM23-positive mitochondria in transduced d*Mfn*KO MEFs. SNVs differ in their capacity to induce the formation of mitochondrial filaments. Overlay images depict differences in the intramitochondrial distributions of MFN2 and TIM23. Bar 10 µm.

### Computational VEP tools based on the analysis of protein features can predict the effect of SNVs on the fusion capacity of MFN2

The large number of MFN2 SNVs subjected and characterized with the cell-based functional assay prompted us to conduct an appraisal of bioinformatic ‘Variant Effect Prediction (VEP)’ tools, *i.e.* tools predicting the functional impact and the pathogenicity of SNVs. The VEP tools currently available can be classified in three main groups according to the data analyzed for prediction of functional impact or pathogenicity (Dong et al., 2015; Katsonis et al., 2022): (1) VEP tools analyzing protein sequence, features, and conservation (*e.g.* SIFT, PolyPhen-2, EVE), (2) VEP tools analyzing nucleotide sequence conservation (*e.g.* PhyloP, SiPhy and GERP++), and (3) VEP tools that score, combine, and integrate the data from different VEP tools and databases (*e.g.* CADD, REVEL, FATHMM, BayesDel and MetaLR). To conduct an unbiased comparison, we analyzed all SNVs with a variety of established VEP tools (supplementary Table 1) belonging to each group and found that different VEP tools provided highly variable predictions for a same mutation (Table 2).

**Table 2:**
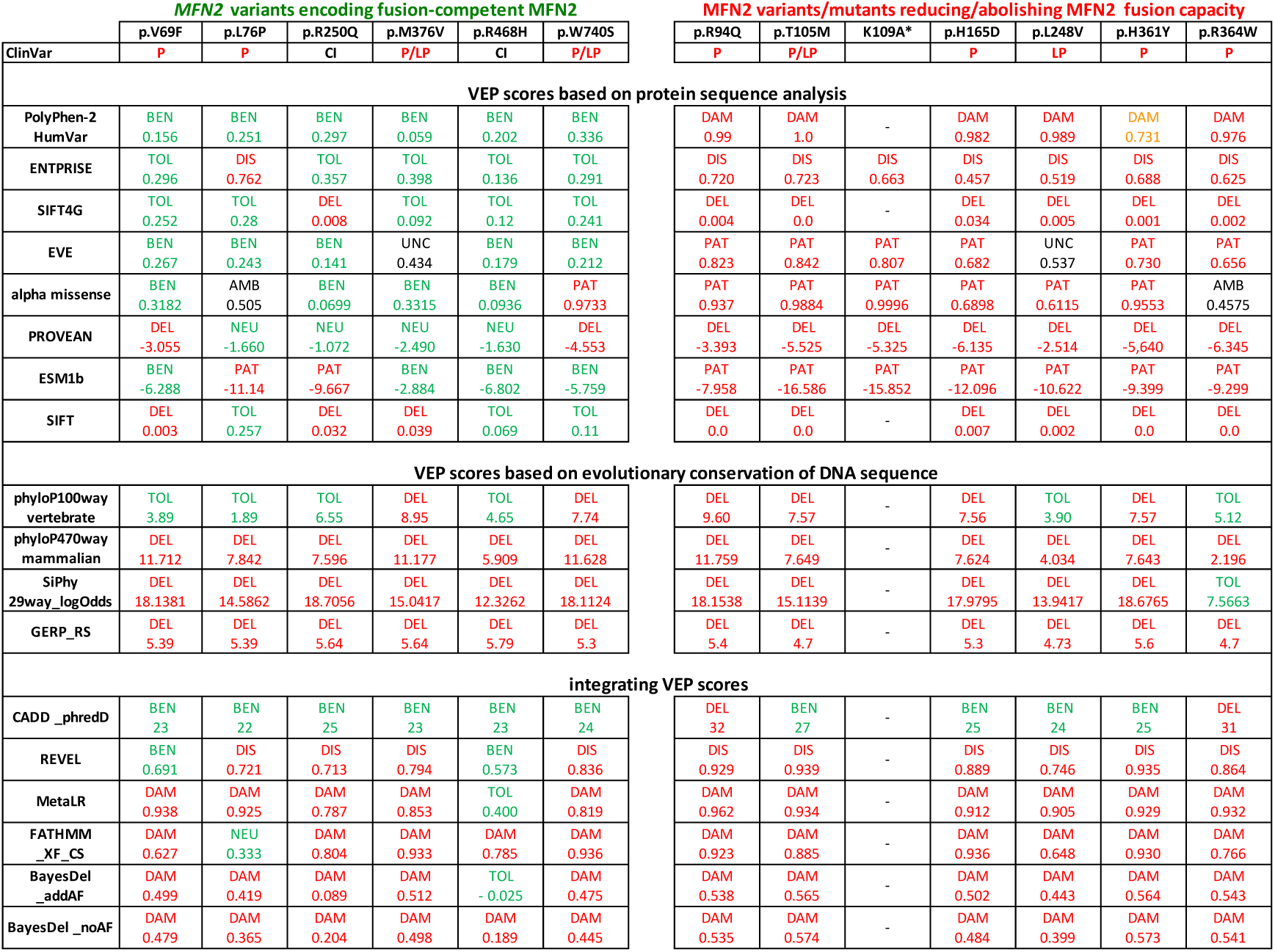
Variant effect predictions (VEPs) based on protein sequence analysis correlate with results of cellular fusion assay. SNVs carrying the indicated amino acid changes were classified according to their impact on the measured fusion capacity. VEP tools were classified according to the analysis strategy: analysis of protein sequence alterations, analysis of DNA conservation and integrative analysis of several parameters and VEP scores. The VEP scores and the resulting interpretation/prediction is indicated. Green: BEN/benign, TOL/tolerated, NEU/neutral. Red: P/pathogenic, LP/likely pathogenic, DAM/damaging, DIS/disease, DEL/deleterious, PAT/pathogenic. Black: CI/conflicting interpretations, UNC/uncertain, AMB/ambiguous. Predictions of protein-based VEP tools correlate with the results of the fusion assay. No correlation is observed with VEP tools based on DNA-sequence analysis or integration of several VEPs.

Interestingly, we discovered that the functional prediction from VEP tools based on the analysis of the protein sequence remarkably matched the results obtained with our cell-based functional test (Table 2). Of note, the occurrence of discordant predictions was very low for four of them (Table 2: PolyPhen-2, ENTPRISE, SIFTG4 and EVE). In contrast, the scores provided by VEP tools based on the analysis of nucleotide sequence conservation (PhyloP, SiPhy and GERP++), as well as the predictions provided by several integrating VEP tools, did not correlate with the experimentally determined fusion capacity of MFN2 (Table 2). In summary, the analysis of functionally characterized SNVs with several established VEP tools indicate that VEP tools based on the analysis of protein sequence (notably PolyPhen-2, ENTPRISE, SIFT4G and EVE) appear suited to predict whether a given SNV affects the fusion capacity of MFN2.

## Discussion

This work describes the development and validation of a mitochondrial fusion assay based on the transduction of human *MFN2* SNVs into OMM-fusion incompetent MEFs (d*Mfn*KO MEFs). This isogenic disease model was chosen as it allows to quantitatively evaluate MFN2 fusogenic activity by performing a basic but straightforward image analysis approach. The robustness of our cell-based OMM-fusion functional test relies on the unprecedented levels of genetic controls introduced to validate SNVs’ sequence integrity after transduction alongside the quantitative analysis of *MFN2* SNVs expression levels using two independent antibodies. Systematic and standardized functional analysis of human *MFN2* SNVs in an isogenic background allowed to classify 12 different *MFN2* SNVs according to their fusion capacity. Furthermore, our experimental strategy also established that the SNVs characterized in this study do not impact the stability or the mitochondrial targeting of MFN2.

In this work, we have established that five out of the 12 mutations subjected to the cell-based OMM-fusion functional test were causing the loss of the fusion capacity of MFN2 (p.R94Q, p.T105M, p.H165D, p.L248V, p.H361Y). Among them, p.R94Q, p.T105M, and p.H361Y had been shown to provoke neurological defects when knocked-in into mice or rats (De Gioia et al., 2020; Sato-Yamada et al., 2022). In contrast, the functional consequence of the two mutations p.H165D and p.L248V had never been investigated since their original identification in patients (Table 1 and (Stuppia et al., 2015)). We further show that human p.R364W induces a reduction of the fusion capacity, as previous reported (Fissi et al., 2020). The low fusion capacity of human p.R364W expressed in MEFs contrasts with the gain of function observed upon expression of its drosophila mimic (*marf*-R404W = MFN2-R364W^like^) in drosophila neurons (Fissi et al., 2018). We hypothesize that this functional divergence could be explained by the large phylogenetic distance and the lower sequence identity between human MFN2 and MARF; however, we cannot fully exclude that such differences might arise from the use of different expression systems and cellular models.

On the other hand, our functional analysis identified six *MFN2* SNVs that were not significantly affecting MFN2 mediated OMM-fusion (Figure 2). The preserved capacity of p.V69F, p.L76P, and p.W740S to mediate OMM-fusion is in agreement with previous functional analysis performed with murine *Mfn2* (Detmer and Chan, 2007). The other three SNVs encoding fusion-competent MFN2 (p.R250Q, p.M376V, and p.R468H) were never functionally characterized since their identification in patients. Noteworthy, none of them provoked an increase in the fusion capacity of MFN2, a gain of function phenotype observed upon expression of variants of *MARF* – the drosophila homologue of MFN2 – in fly neurons (Fissi et al., 2018).

Sequence analysis revealed that most of the loss of function mutations are localized within (p.T105M, p.H165D, p.L248V) or in close vicinity (p.R94Q, p.H361Y, p.R364W) to the GTPase domain (Figure 1A). To gain further insight into the localization of mutations, we visualized them on the 3D-structure of functional domains of MFN1 and MFN2, as well as of BDLP, a bacterial MFN-homologue (Low and Löwe, 2006; Low et al., 2009; Yan et al., 2018; Li et al., 2019) (supplementary Figure 3). Interestingly, residues affecting the fusion capacity of MFN2 located to the GTP-binding pocket (p.T105M, p.K109A and p.H165D) or to a region of the GTP-binding domain that contacts the four-helix bundle in the GTP-bound conformation (p.R94Q, p.L248V, p.H361Y and p.R364W). This likely indicates that these SNVs may provoke a decrease or loss of MFN2’s fusion capacity by altering GTP-binding or hydrolysis, as well as associated conformational changes.

Comparison of the functional classification of *MFN2* SNVs with computational predictions obtained from 16 ‘variant effect prediction (VEP)’ tools unraveled that predictions with tools based on the analysis of protein sequence correlated remarkably with the results of our functional analyses (Table 2). In fact, a majority of *MFN2* SNVs retaining their OMM-fusion capacity were predicted to be tolerated or benign whereas, conversely, a majority of *MFN2* SNVs causing loss of function were predicted to be damaging or pathogenic (Table 2). In contrast, the correlation with other VEP tools was poor (Table 2).

It is important to stress out that our functional test, specifically determining the impact of SNVs on the OMM-fusion activity of MFN2, can support the pathogenic character of fusion-incompetent *MFN2* mutants. In contrast, the absence of a defect does not necessarily imply that corresponding SNVs can be classified as benign. Indeed, such SNVs maybe benign (*e.g.* p.R468H) or affect other functions of MFN2 escaping, at this stage, the perimeter of our functional assay: mitochondrial transport (Misko et al., 2012; Baloh et al., 2007), ER-mitochondrial tethering (De Brito and Scorrano, 2008) (Filadi et al., 2015) or autophagy (Zhao et al., 2012) (Chen and Dorn, 2013). Alternatively, it can be envisioned that this cellular model minors the fusion or other MFN2-associated phenotypes which would be more efficiently unmasked in neurons (*e.g.* mitochondrial mobility along axons and dendrites). For instance, the p.V69F, p.L76P, and p.W740S variants (identified by positional cloning (Züchner et al., 2004)), were shown to alter mitochondrial mobility in cultured neurons (Baloh et al., 2007).

To conclude, we believe that the development of an innovative mitochondrial fusion assay, the screening of *MFN2* VUS and the identification of VEP tools able to predict fusion defects represent solid findings and potent tools for the interpretation and classification of *MFN2* variants identified in CMT2A and in other neuropathies. Our future goal is to improve the sensitivity and resolution of the cell-based functional assay and to widen the scope of this test by investigating further *MFN2* variants, other MFN2 activities as well as developing a neuronal isogenic CMT2A cell model. Beyond supporting diagnosis of CMT2A and of CMT-related neuropathies, this work will improve our knowledge of MFN2 functions and broaden our understanding of pathological mechanisms involved in CMT2A.

## Materials and Methods

### Databases and Bioinformatics Analysis

The SNVs of *MFN2* linked to CMT2A (Table 1) were identified and selected in ClinVar, HGMD and Inherited Neuropathy Variant Browser (Landrum et al., 2020) (Stenson et al., 2014) (Saghira et al., 2018) (supplementary Table 1) as well as in a review by Pipis and co-workers (Pipis et al., 2020). The tools for ‘Variant Effect Prediction (VEP)’ as well as the websites where the analysis was performed and/or the predictions downloaded are indicated in supplementary Table 1 (Fokkema et al., 2021; Karczewski et al., 2020; McLaren et al., 2016; Liu et al., 2011; 2020; Adzhubei et al., 2010; Frazer et al., 2021; Cheng et al., 2023; Brandes et al., 2023; Kumar et al., 2009; Vaser et al., 2016; Hongyi Zhou, 2016; Choi et al., 2012; Pollard et al., 2010; Garber et al., 2009; Davydov et al., 2010; Rentzsch et al., 2019; Ioannidis et al., 2016; Dong et al., 2015; Shihab et al., 2013; Feng, 2017). Unless otherwise indicated, predictions rely on the default threshold values. For ESM1B, we applied a threshold (−7.5) yielding a true-positive rate of 81% and a true-negative rate of 82% in both datasets (Brandes et al., 2023). We applied the threshold values proposed by Dong et al. (Dong et al., 2015) for GERP++RS (>4.4), Siphy (>12.17) and phyloP470way mammalian (>1.6), and a threshold value equivalent to purifying selection by ‘loss of function’ (> 7.5) for phyloP100way vertebrate (Vy et al., 2021).

### Cloning and mutagenesis

Variants of human *MFN2* were generated by mutagenesis using a QuickChange-derived protocol (Xia et al., 2015). The cDNA encoding wild-type human *MFN2* (transcript variant 1, accession NM_014874.4 (Rojo et al., 2002)) was either mutagenized in a 3 kb cloning plasmid (pKSPS (Bahri et al., 2021)) before subcloning into pQCXIB (the retroviral expression vector, Addgene plasmid #22800) or was directly mutagenized in pQCXIB. For convenience, the p.L76P variant was mimicked by the change of two nucleotides (ctg>ccc) instead of one (ctg>ccg). The sequence of *MFN2* variants was verified by sequencing using Mix2Seq kits from Eurofins Genomics. All plasmids will be made available via a plasmid repository.

### Cell Culture and transduction

Mouse embryonic fibroblasts (MEFS) were cultured in Dulbecco’s Modified Eagle’s Medium (DMEM) containing 1 mM sodium pyruvate and 4.5 g/L Glucose (Dutscher, L0106-500), supplemented with 7% fetal bovine serum (FBS) (PAN-Biotech – P30-3306), 2mM glutamine (Dutscher – X0551-100) and 1% penicillin/streptomycin (PAN-Biotech - P06-07100), at 37 °C in an incubator with a humidified atmosphere of 5% CO_2_. Reaching 80% confluency, cells were passaged by trypsinization using Trypsin-EDTA (PAN-Biotech - P10-023100) and were allowed to adhere and grow for 36 to 48 hours before sample collection for analysis. Two different cultures of wild-type MEFs were used as controls: CTRL_A_ (*Opa1*^+/+^ (Silva Ramos et al., 2019)) and CTRL_B_ (*Atg5*^+/+^ (Frank et al., 2012)).

Double *Mfn1/Mfn2* knock-out (d*Mfn*KO) MEFs (Koshiba et al., 2004) expressing variants of human MFN2 were generated by stable transduction as described in el Fissi et al. (Fissi et al., 2018). Essentially, viral particles were generated by transfection of Plat-E retroviral packaging cells ((Morita et al., 2000) Cell Biolabs, Inc.) with retroviral pQCXIB-plasmids encoding the indicated *MFN2* variants. MEFs were transduced with viral supernatants diluted in complete culture medium and supplemented with 8 µg/ml polybrene and 5 µg/ml of plasmocin; transduced cells were then selected by addition of blasticidin at a final concentration of 20 µg/ml. To ensure transduction of a majority of cells with a single vector copy, viral supernatants were diluted to achieve transduction efficiencies below 50% (Fehse et al., 2004).

### Sequencing of human *MFN2* variants transduced into MEF’s genomes

Cells were grown on 100 mm diameter culture Petri dishes for 48 hours, washed with PBS before trypsinization, and collected as cell pellets by centrifugation. DNA extraction was then completed using the DNeasy Tissue and Blood Kit (Qiagen, 69504) following the manufacturer’s instructions. DNA was later quantified using the Helixyte Green^TM^ dsDNA Quantification Kit Green Fluorescence (AAT Bioquest, 17651).

Standard PCR was then completed using Phusion™ High-Fidelity DNA Polymerase (Thermofisher, F530XL) in order to amplify fragments of the *MFN2* variants’ cDNA integrated into the MEF genome by retroviral transduction. Four different primer couples (supplementary Table 2) were used in order to generate overlapping fragments covering the entirety of MFN2. Cycling conditions were as follows: 98°C for 5 min for one cycle, then 98°C for 10 seconds followed by annealing at 60°C for 30 seconds, and 72°C for 3 minutes for 40 cycles, and 72°C for 7 minutes as a final cycle.

The PCR products were then migrated on a 1.2% agarose gel supplemented with SYBR Safe DNA gel Stain (Invitrogen, S33102) and DNA was then purified from the obtained bands using the GeneJET Gel Extraction Kit (Thermoscientific, K0692) following the manufacturer’s instructions. The samples were then sequenced using Mix2Seq kits from Eurofins Genomics.

### Western Blot

Cells were grown on 100 mm diameter culture Petri dishes for 48 hours, then washed with PBS twice before harvesting the adherent fibroblasts by scraping in ice cold PBS. Cells in suspension were then pelleted by centrifugation (1,000g for 10 minutes) and frozen after removal of the PBS supernatant. Protein lysis was then achieved by resuspending the −80°C stored pellets in 50µL of RIPA consisting of 50 mM Tris pH 8, 5 mM EDTA pH 8, 150 mM sodium chloride, 1 % NP-40, and 0.5 % sodium deoxycholate, 0.1 % SDS for 15 min on ice; the latter lysis buffer was supplemented with cOmplete™, Mini, EDTA-free Protease Inhibitor Cocktail (Roche –11836170001).

Protein concentration in the lysates was determined with the DC Protein Assay Kit (Biorad – 5000113 to 5000115) using BSA as a standard. Loading samples were prepared in Laemmli sample buffer containing 390 mM thioglycerol at a final protein concentration of 1.5 µg/µL. After heating the samples for 5 min at 95°C, a total of 35µg of protein was resolved on 10% Tris-glycine polyacrylamide gel and then transferred to 0,2 µm nitrocellulose membranes by wet transfer for 1 hour at 90V.

Membranes were subsequently blocked for 20 min in 5% Non-Fat dry milk powder in TBS-Tween (TBS + 0,05% Tween 20) after Ponceau red staining and colorimetric image capturing. Immunoblotting was next performed by incubating the membranes overnight at 4°C with the following primary antibodies: mouse monoclonal anti-mitofusin 2 antibody [6A8] (abcam, ab56889, dilution of 1/1000), rabbit polyclonal anti-mitofusin 2 (abcam, ab50838, dilution of 1/1000), and mouse monoclonal anti-beta actin (ProteinBiotech, 66009-1, dilution of 1/30000). The following species-specific HRP-conjugated secondary antibodies (diluted in 3% Non-Fat dry milk powder) were used: Peroxidase AffiniPure Goat Anti-Mouse IgG (H+L) (Jackson Immuno Research, 115-035-062, dilution of 1/10000), and Peroxidase AffiniPure Goat Anti-Rabbit IgG (H+L) (Jackson Immuno Research, 115-035-003, dilution of 1/10000) and the membranes were incubated for 1 hour at room temperature in the respective HRP-conjugated antibodies. Following two washes in TBS-Tween and a final wash in TBS, ECL detection was performed using the Clarity Western ECL Substrate kit from Biorad (170-5061) and chemiluminescence imaging was completed with an Amersham™ ImageQuant 800 Fluor. To note, for consecutive decorations with rabbit or mouse antibodies, the HRP-signal of the first antibody was a deactivated by incubation of the membranes with a 0.1% sodium azide solution for 30 min. For quantitative analysis, the intensity of the signal was determined using ImageJ software; MFN2 signal was normalized to d*Mfn*KO+MFN2 and to beta actin in blots of four independent experiments. The relative protein expression levels obtained were analyzed by one-way ANOVA with Kruskal-Wallis Multiple Comparison post hoc test. *P < 0.05, **P < 0.01, ***P < 0.01 compared with d*Mfn*KO + MFN2_A_.

### Immunofluorescence microscopy and analysis of mitochondrial morphology

Cells plated onto glass coverslips were fixed in 3.2% paraformaldehyde for 20 min at room temperature, then washed once with PBS before being permeabilized using PBS with 0.1% Triton X-100 (PBST) solution for 5 min.

In order to study the mitochondrial morphology of the cells, cells were treated with a 8M urea solution for 20 min (Malka et al., 2007) and decorated for 1h30min with mouse monoclonal anti-VDAC1/Porin + VDAC3 antibody [20B12AF2] (abcam, ab14734, dilution of 1/400) as a mitochondrial marker. To determine the localization of MFN2, the following procedure was completed. After 30 minutes of blocking with a 10% BSA solution, the coverslips were decorated with the following primary antibodies diluted in a 3% BSA solution for 1h30min at room temperature: mouse monoclonal anti-TIM23 (BD Transduction Laboratories, 611223, dilution of 1/400) and rabbit monoclonal anti-Mitofusin-2 (D1E9) (Cell Signaling, 11925, dilution of 1/100).

Incubation with the following secondary antibodies Goat anti-Mouse IgG (H+L) Alexa Fluor™ Plus 488 (Invitrogen, A32723, dilution of 1/800) and Goat anti-Rabbit IgG (H+L) Alexa Fluor™ Plus 555 (Invitrogen, A32732, dilution of 1/800) was then completed for 45 min at room temperature following a quick wash with PBST. Finally, the coverslips were washed with PBST, then PBS, and distilled water before being mounted with Mowiol mounting medium supplemented with 0.5 µg/ml DAPI (Malka et al., 2007).

Images were acquired using an inverted Microscope Olympus (Olympus IX81) with 60X and 100X oil objectives and were analyzed and processed using Olympus cellSens and ImageJ software. For quantitative analysis (Figure 2 and supplementary Figure 1), mitochondrial morphology of 150–200 transfected cells was determined in three independent experiments as filamentous (FIL, with a dense network of elongated and/or interconnected filaments) or fragmented (FRA, lacking filaments and displaying separate punctate or round mitochondria). Cells that did not fit into any of these categories were classified as intermediate (INT, with a mixture of punctate mitochondria and few or short mitochondrial filaments) (Supplementary Figure1). A two-way ANOVA statistical analysis with Dunnett’s Multiple Comparison post hoc test was conducted using Prism (GraphPad) *P < 0.05, **P < 0.01, ***P < 0.01 compared with the d*Mfn*KO MEFs.

## Acknowledgements

We thank David Chan, Stéphane Duvezin-Caubet and Eric Campeau for providing d*Mfn*KO MEFs, ATG5+/+ MEFs and pQCXIB plasmid (Addgene plasmid #22800), respectively. We are grateful to Jim Dompierre for advice and assistance with fluorescence microscopy, to Esra Karatas and Anaïs Rey for help with the transduction and characterization of MEFs and to Macha Nikolski for assistance with databases. We are indebted to Tanja Stojkovic, Julian Cassereau, Guilhem Solé, Phillipe Pasdois and Nathalie Bernard-Marissal for fruitful discussions and valuable comments on the manuscript. We express our gratitude to Eulalie Lasseaux for sharing her expertise and insights on the use and interpretation of Variant Effect Predictors. This work was supported by grants from Bordeaux University (GPR Light), AFM (24219), and ANR (ANR-16-CE14-0013 and ANR-22-CE14-0040).

## Supplementary Tables and Figures for Manuscript

**Supplementary table 1:**
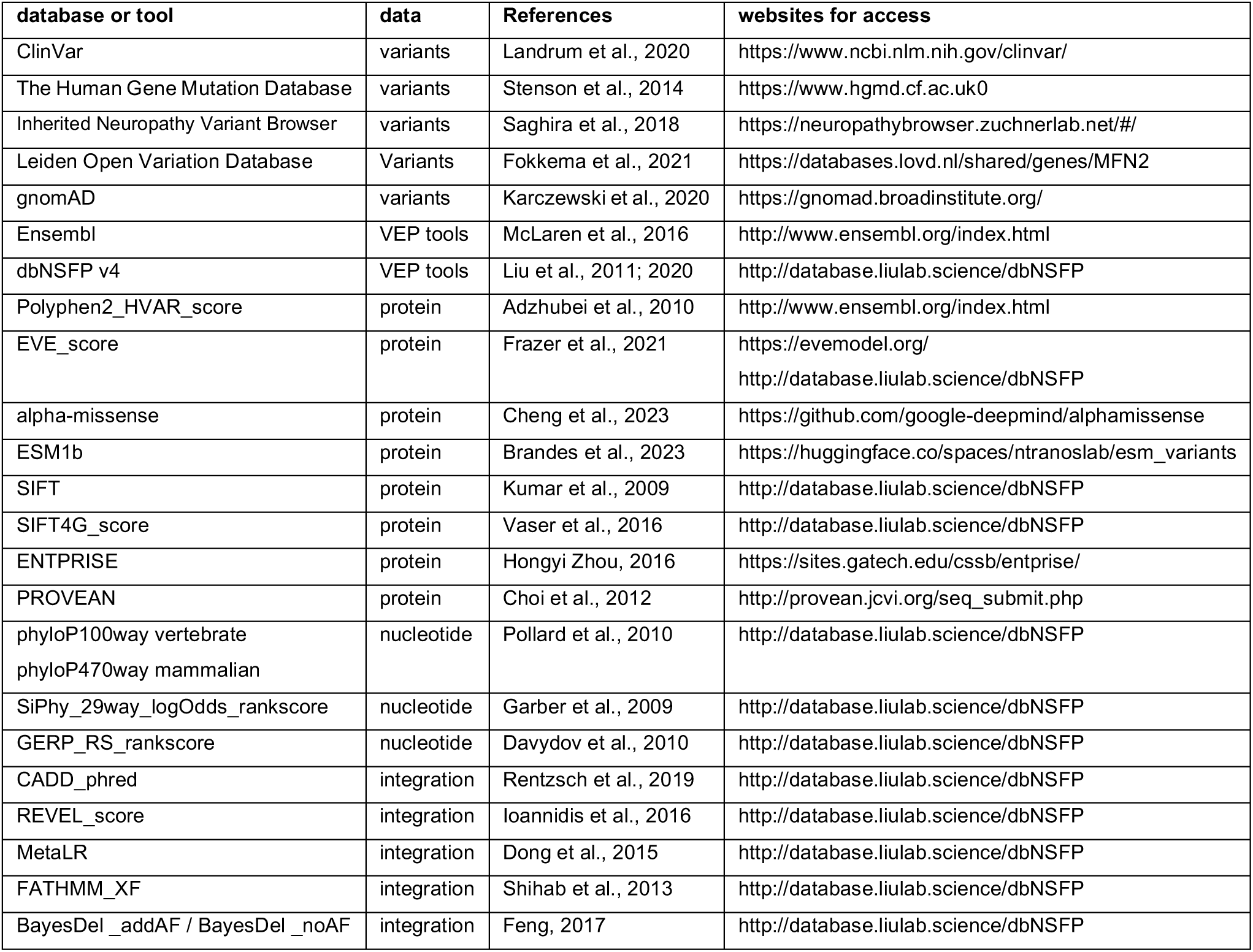
Databases and websites hosting clinical, genetic and genomic data (ClinVar, HGMD, gnomAD) or Variant Effect Predictor (VEP) tools (Ensembl, dbNSFP v4). The data column indicates the data analysed by VEP tools to predict conservation, dysfunction and/or pathogenicity. Integration indicates integration of clinical findings and/or the predictions of various VEP tools.

**Supplementary table 2.**
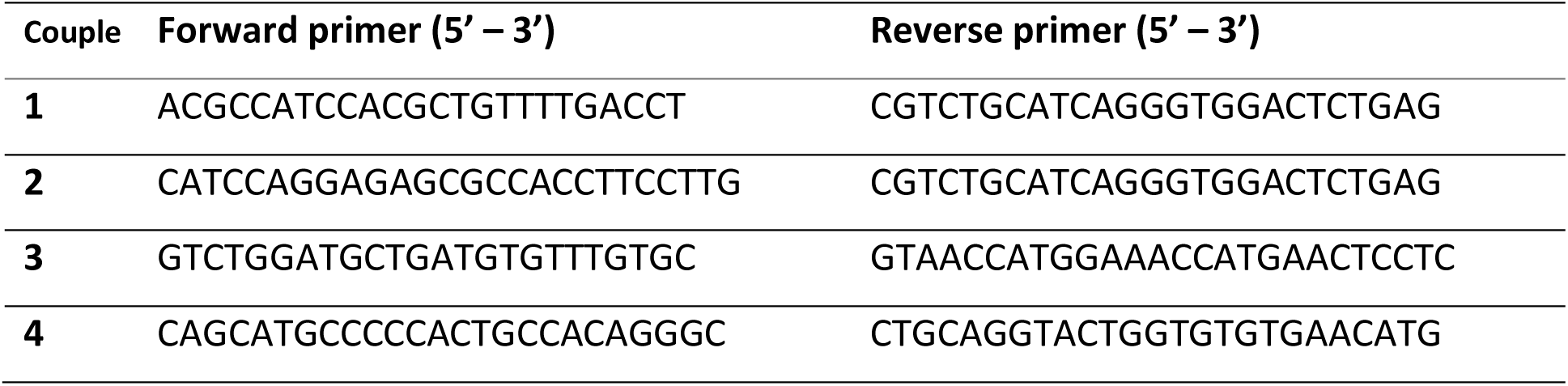
Oligonucleotides for amplification and sequencing of MFN2 cDNA.

**Supplementary Figure 1.**
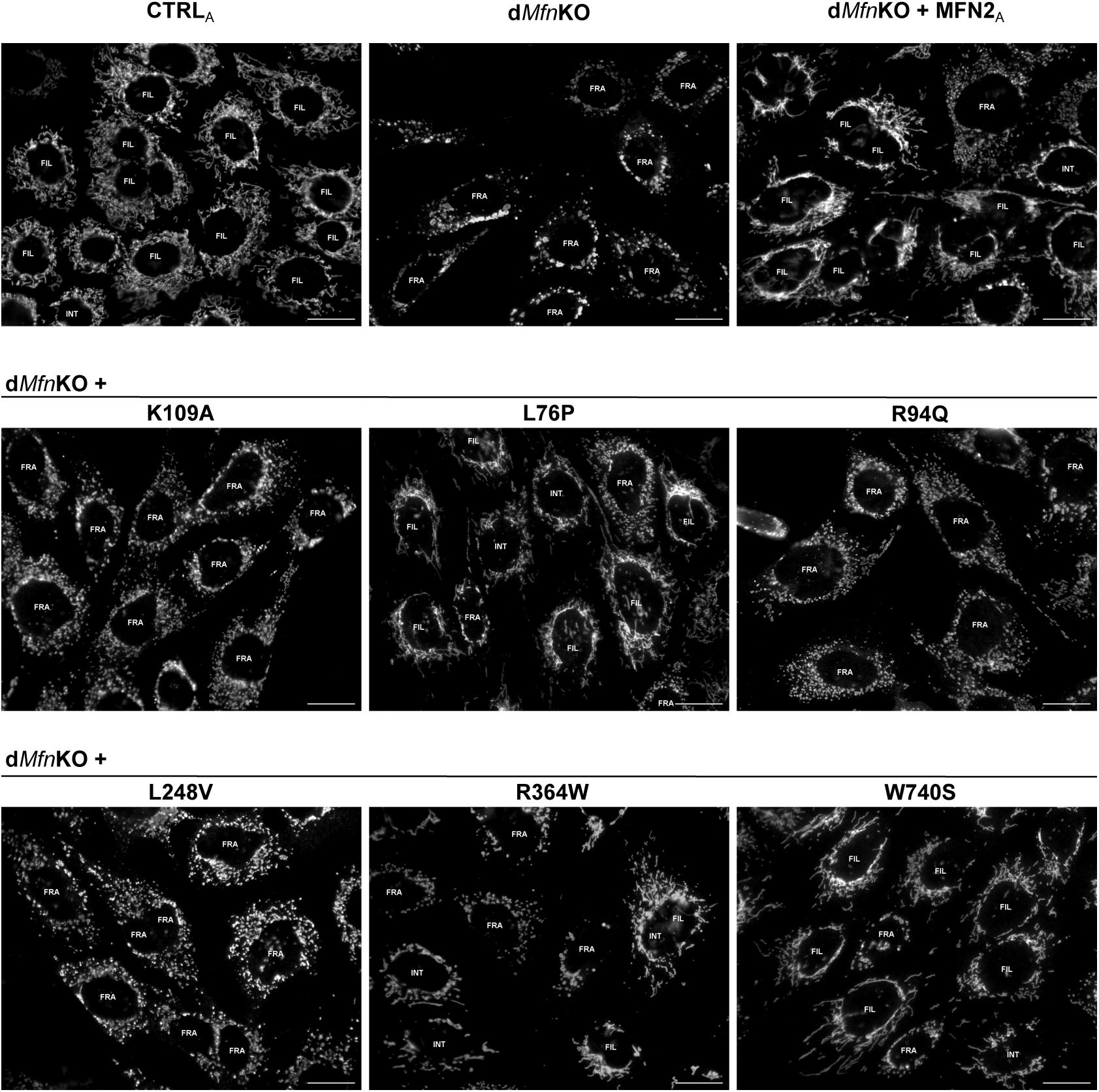
Visualization and assessment of mitochondrial morphology by immufluorescence microscopy. Representative immunofluorescence images of wild-type MEFs (CTRL_A_), untransduced d*Mfn*KO MEFs and d*Mfn*KO MEFs transduced with wild-type *MFN2* (MFN2_A_) or the indicated variants. The overall mitochondrial morphology of cells stained with antibodies against the mitochondrial marker VDAC was classified as fragmented (FRA), filamentous (FIL) or intermediate (INT). Bar: 20 µm.

**Supplementary Figure 2.**
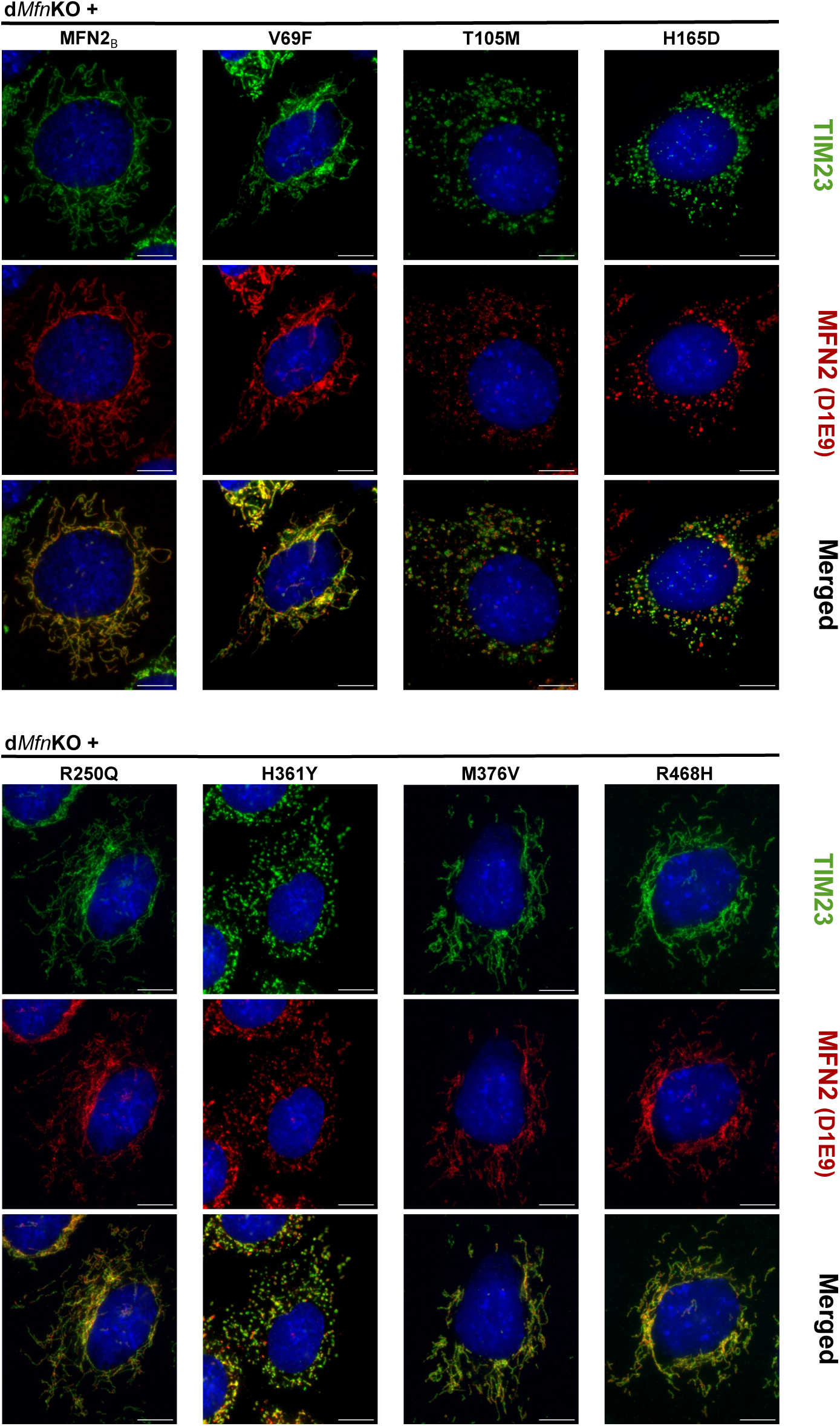
Mitochondrial localization of human MFN2 variants expressed in d*Mfn*KO MEFs. Representative immunofluorescence images of cells stained with antibodies against the mitochondrial marker TIM23 (green) and MFN2 (D1E9, red) and with the nuclear stain DAPI (blue). MFN2, undetectable in untransduced d*Mfn*KO MEFs, localizes to TIM23-positive mitochondria in transduced d*Mfn*KO MEFs. MFN2 variants differ in their capacity to induce the formation of mitochondrial filaments. Bar: 10 µm.

**Supplementary Figure 3.**
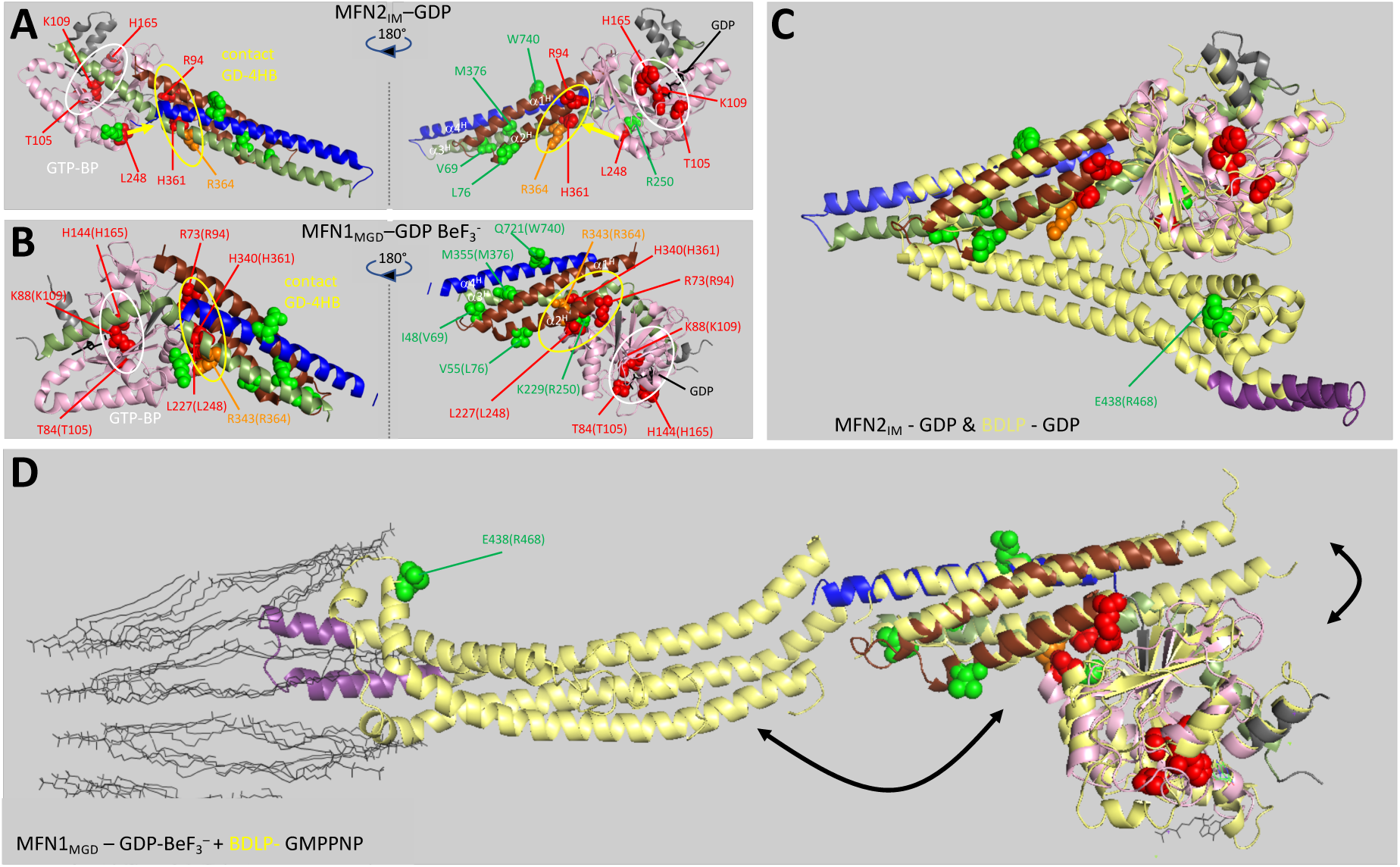
Relative position of the amino acids altered by *MFN2* variants within the 3D structure of MFN1, MFN2 or BDLP. The position of MFN2 amino acids or their homologs is indicated. The color code corresponds to the fusion capacity of protein variants: red (abolished), orange (diminished) or green (not or slightly affected). **A**: ClustalW alignment of human mitofusins MFN1 and MFN2. Functional domains are underlined in light grey homologous aminoacids in yellow. Color code of alignment: red/identical, green/highly similar, blue/weakly similar, black/different amino acids. **A/B**: Structure of a minimal functional domain of MFN2 (Mfn2_IM_) bound to GDP (**A**) or of MFN1 (MFN1_MGD_) bound to GDP BeF_3_^-^, a GTP mimick (**B**). The positions of MFN2 protein variants (**A**) or of MFN1 residues homologous to MFN2 protein variants (**B**, in brackets) are indicated. The GTP-binding domain is colored in pink. The alpha-helices of the four-helix bundle (α1^H^, α2^H^, α3^H^ and α4^H^) are colored in brown, green and blue, respectively. The GTP-binding pocket (GTP-BP) and the contact area between GTP-binding and four-helix-bundle (GD-4HB) are indicated in white and yellow, respectively. **C/D:** Structural alignment of MFN2_IM_ or MFN1_MGD_ (colored as in **A/B**) with a bacterial MFN-homologue (BDLP) bound to GDP (**C**) or GMPPNP, a non-hydrolysable GTP-analogue (**D**). BDLP is colored in yellow but for the domain mediating membrane-anchoring (magenta). The phospholipids of a model membrane are shown in dark grey. Black arrows point to differences between the different, nucleotide-induced conformations.

### PDB accession numbers

6jfk.pdb (MFN2 GDP), 5yew.pdb (MFN1 GDP BeF_3_^-^), 2j68.pbb (BLDP GDP), 2w6d.pdb (BDLP GMPPNP).

